# Testosterone secretion varies in a sex- and stage-specific manner: insights on the regulation of competitive traits from a sex-role reversed species

**DOI:** 10.1101/775262

**Authors:** Sara E. Lipshutz, Kimberly A. Rosvall

## Abstract

Testosterone (T) mediates a variety of traits that function in competition for mates, including territorial aggression, ornaments, armaments, and gametogenesis. The link between T and mating competition has been studied mainly in males, but females also face selection pressures to compete for mates. Sex-role reversed species, in which females are the more competitive sex, provide a unique perspective on the role of T in promoting competitive traits. Here, we examine patterns of T secretion in sex-role reversed Northern Jacanas (*Jacana spinosa*) during breeding, when females are fertile and males are either seeking copulations or conducting parental care. We measured baseline levels of T in circulation along with a suite of behavioral and morphological traits putatively involved in mating competition. We evaluated hypotheses that levels of T track gonadal sex and parental role, and we begin to investigate whether T and competitive traits co-vary in a sex- and stage-specific manner. Although females had higher expression of competitive traits than males at either breeding stage, we found that females and incubating males had similar levels of T secretion, which were lower than those observed in copulating males. T was correlated with wing spur length in females and testes mass in copulating males, but was otherwise uncorrelated with other competitive traits. These findings suggest that levels of T in circulation alone do not predict variation in competitive traits across levels of analysis, including gonadal sex and parental role. Instead, our findings coupled with prior research indicate that selection for female mating competition and male care may generate different physiological regulation of competitive traits in jacanas.

**Highlights:**

- In role reversed species, females face stronger competition and males care for offspring
- We examine testosterone (T) and competitive traits in female and male jacanas
- Circulating T is similar in females and incubating males, but higher in copulating males
- Both gonadal sex and parental role shape patterns of T secretion
- Physiological regulation of competitive traits may differ in female and male jacanas

## 1. Introduction

For males of many vertebrate species, testosterone (T) has been associated with phenotypic traits that lead to successful competition over mates. These traits include aggressive behavior (Wingfield et al., 1990), weaponry (Malo et al., 2009), body size (Cox et al., 2009), ornamentation (McGlothlin et al., 2008), and gonadal size (Preston et al., 2012). Females also express many of these same traits, which likewise function in competition for mates and other breeding resources (Clutton-Brock, 2009; Hare and Simmons, 2018; Rosvall, 2013a; Tobias et al., 2012). Like males, females secrete T and have the physiological machinery to respond to T (Staub and De Beer, 1997). However, the relationship between T and mating competition in females has found more equivocal support (Goymann and Wingfield, 2014; Rosvall et al., in press, Cain and Ketterson 2012), suggesting that T may influence trait expression differently in males and females. This can be explained in part by sex differences in the selective pressures that drive endocrine mechanisms of behavior (Wingfield et al. 1990). For instance, the relative importance of aggression versus parental care in males and females may shape both sexual dimorphism in T, as well behavioral sensitivity to T (Lynn, 2008; Rosvall, 2013b) – this is one hypothesized driver for why females tend to have lower levels of T in circulation than males.

Levels of T in circulation are typically elevated at the beginning of the breeding season, when competition for mating opportunities is high, and then T levels decline as behavioral efforts shift to parental care (Wingfield et al. 1990). This cross-stage shift in T production has been well demonstrated in males (Hirschenhauser and Oliveira, 2006), and is potentially more dramatic in females (DeVries et al., 2012; George and Rosvall, 2018; Jawor et al., 2007). Cross-stage shifts may also alter correlations between T and sexually selected traits, such that these traits are strongly integrated with T during competition for mates, but more independent from T during periods of parental care (Ketterson et al., 2009; Lipshutz et al., 2019a).

In sex-role reversed species, females face stronger selection to compete for mates, and males predominantly care for offspring (Emlen and Oring, 1977), providing a unique opportunity to disentangle parental role from gonadal sex as drivers of variation in T. Whereas sex-role reversed females are freed from the constraints of parental care, selective pressures to care for offspring may limit T levels in males. The hypothesis that sex-role reversed females have higher T than males has intrigued behavioral ecologists for decades. Whereas early studies in sex-role reversed phalaropes found higher T in ovarian than testicular tissue (Höhn, 1970; Höhn and Cheng, 1967), most studies have found that the direction and magnitude of sex differences in T is typically similar between role-reversed species and species with traditional sex roles (Wilson’s phalaropes - Fivizzani et al., 1986; red-necked phalarope - Gratto-Trevor et al., 1990; spotted sandpiper - Rissman and Wingfield, 1984, African black coucal - Goymann et al., 2004; but see barred button-quails Muck and Goymann 2011). Approaching sex-role reversed species from a hormonal perspective can shed new light on the regulation of competitive traits, which may vary based on gonadal sex and/or parental role.

One of the most well-known examples of sex-role reversal is the jacana (family Jacanidae). Although both sexes aggressively compete for territories, females face more intense sexual selection, have higher potential reproductive rates, and have female-biased operational sex ratios (Emlen and Wrege, 2004a). Female jacanas also have larger secondary sexual traits, including weaponry, body mass, ornamentation, and behavioral dominance (Emlen and Wrege, 2004a; Lipshutz, 2017; Stephens, 1984). Males provide nearly all parental care, which includes incubation and foraging with chicks (Emlen and Wrege, 2004b; Jenni and Collier, 1972). Although jacanas are a classic example of female-biased dimorphism in behavior and morphology (Emlen and Oring, 1977), there is currently no published work on the physiological mechanisms that regulate the expression of these traits, stemming in part from the logistical challenges of research on these tropical birds in their remote wetland habitats.

Here, we quantified circulating T levels in free-living, sex-role reversed jacanas to evaluate the hypotheses that levels of T track gonadal sex and/or parental role. Sex-role reversed females should be unconstrained by parental care, and if they have higher T than males, this would suggest that T levels are shaped by parental roles more so than gonadal sex. We further predict that males incubating eggs should have lower T than males seeking copulations. As a secondary goal, we begin to evaluate whether co-variation between T and competitive traits varies in a sex- and breeding stage-specific manner. Few studies have examined phenotypic co-variation with T in sex-role reversed species, but one found that T positively correlated with plumage coloration and body condition in females only (Muck and Goymann, 2011), and the other found that ornaments were testosterone-dependent in both sexes (Eens et al., 2000). Neither of these studies examined the influence of breeding stage on phenotypic co-variation, which is important for considering how temporally variable selection pressures (i.e. mating competition vs. parental care) shape differences both between and within the sexes. We predict that T will be more strongly linked with competitive traits in females and copulating males, but that T should be more independent from these traits in incubating males.

## 2. Materials and Methods

### 2.1. Subjects

Northern Jacanas (*Jacana spinosa*) are tropical shorebirds found throughout Central America, from Mexico to Panama. They breed asynchronously and year-round, although breeding increases during the rainy season in Panama from roughly May to October. We conducted fieldwork from 4 June to 9 July 2018 in La Barqueta, Chiriqui (8.207N, 82.579W).

### 2.2. Aggression assays, plasma collection, and morphological measurements

Prior to each aggression assay, we observed each individual for several days to ensure that it was pair-bonded (e.g. foraging with mate) and territorial (e.g. actively defending territory from intruding floaters). Territorial residents behave distinctly from floaters, who do not breed nor defend territories (Emlen and Wrege, 2004a). We also determined breeding status based on whether males were copulating or incubating a nest. We did not color-band individuals as in previous studies (Lipshutz et al., 2019b) and were therefore unable to determine harem size, but territory holders have high site fidelity (Emlen and Wrege, 2004a), and the agricultural land where we conducted this research has small ponds and canals where territorial boundaries are distinct. We only sampled individuals that we observed in these same stable locations consistently each and every day. We simulated territorial intrusion in males (n = 12) and females (n = 10) using female taxidermic mounts and conspecific vocalizations following Lipshutz (2017). Briefly, we set up a camouflage blind and placed the mount and speaker in the center of a female and one of her male mates’ territory (∼15m from the nest if the male was incubating). We used a 10-sec recording of jacanas fighting and vocalizing to attract the focal individual, and began the assay the moment the female or male responded. Each assay presented a random combination of 4 acoustic stimuli and 4 taxidermic mounts. We measured a suite of aggressive behaviors, including average distance from the mount, hover flights, wing spreads, flyovers, and vocalizations, described previously in an ethogram for jacanas (Lipshutz, 2017). There is some evidence that T may elevate after an experimental intrusion (Wingfield and Wada 1989), but recent analyses suggest that this hormonal response is uncommon in birds (reviewed in Wingfield et al. in press). We nevertheless limited trials to 5 min, and promptly attempted to capture each individual, to obtain unprovoked T levels.

Following the assay, 9 individuals were captured and shortly thereafter euthanized for another study (i.e. “immediate collection”). We collected individuals using an air rifle, followed by an anaesthetic overdose of isoflurane and decapitation to collect trunk blood (average time from assay start to euthanasia = 9 min 20 sec ± 1 min 35 sec for immediate collection individuals). We were unable to collect 13 individuals within 10 minutes post-assay, and so, we returned 5-8 days later to collect these individuals (“i.e. delayed collection”, n = 6 females, 5 incubating males, and 1 copulating male). For delayed collections, we ensured that male breeding stage did not change; for instance, if a male was incubating on the day of the assay, we monitored with daily behavioral observations to confirm that he was still incubating on the day of collection. For each individual, we collected whole blood into heparinized BD Microtainers (product #365965) and stored on an ice pack until we separated plasma by centrifuging for 10 minutes at 10,000 rpm. We stored plasma at −20°C for later testosterone assays.

Males and females were tested independently of each other, except for one pair which was collected immediately after the same aggression assay. For each sex, we balanced immediate and delayed collection sample sizes. We confirmed sex and breeding status by examining whether females had hierarchical follicles, and whether males had brood patches. We determined that 7 males were incubating, 5 were copulating, and all 10 females were fertile, with hierarchical follicles. Sample sizes for males in each breeding stage were smaller than anticipated due to challenges with both locating nests and capturing individuals.

### 2.3. Morphological measurements

Postmortem, we measured several traits putatively involved in mating competition in jacanas: wing spur length, body mass, facial shield length, and gonad mass. In a congener, the Wattled Jacana (*J. jacana*), territorial resident status was associated with larger wing spurs, facial shields, and body mass for both sexes, and only territory holders can breed (Emlen and Wrege, 2004a). These traits are much smaller in floaters, who do not obtain territories or reproduce. We assume that these traits similarly relate to intra-sexual competition in Northern Jacanas, although we have not tested this explicitly. For both species, wing spurs, facial shields, and body mass are significantly larger in adult female jacanas compared to males (Emlen and Wrege, 2004a; Lipshutz, 2017), but these traits are small in juveniles of both sexes (Lipshutz, personal observation). We measured body mass with a digital scale (0.01g), and we measured wing spur length (from base center to tip, 0.1 mm) with calipers. Wing spurs are sharp, yellow keratinous sheaths over metacarpal bone growths that jacanas use as weapons and display during aggressive posturing. In the Wattled Jacana, wing spurs length positively correlated with age in males but not females, perhaps because they were worn down by abrasion from fighting (Emlen and Wrege, 2004a). However, we did not observe any worn down wing spurs in this study. In Northern Jacanas, facial shields are yellow, fleshy, and extend from the upper mandible to the forehead. We measured facial shield length (from right nare to top right lobe, 0.1 mm) with calipers. We also measured gonad mass with a digital gem scale (0.1oz). Gonad mass has clear connections to mating competition because jacanas are polyandrous - males that are simultaneously mated with a single female compete to fertilize her eggs, and females are continuously producing eggs for available mates to incubate, (Emlen et al., 1989). Testes size is associated with T in many avian species (Garamszegi et al., 2005), and testes size is correlated with sperm length in shorebirds (Johnson and Briskie, 1999).

### 2.4. Testosterone enzyme immunoassay

We extracted steroids from plasma samples using diethyl ether (3x extractions) and reconstituted in 250μL assay buffer. We measured testosterone using a High Sensitivity Testosterone ELISA kit (Enzo #ADI-900-176, Farmingdale, NY, USA) following methods described in George and Rosvall (2018). We confirmed assay parallelism by comparing concentrations from a standard curve made by kit standards to a displacement curve made by dilutions of a copulating male jacana (R^2^ = 96.3%). We ran all samples in duplicate. We initially used 40μL plasma from females, and 20μL from males. Samples from three females and four copulating males initially showed less than 20% maximum binding, and two males showed greater than 80% maximum binding, so we re-ran them using 10μL plasma and 40 uL of plasma, respectively, to obtain values in the most sensitive part of the curve. We calculated T concentration by comparing sample absorbance with the absorbance of the assay’s standard curve (Gen5 curve-fitting software, Biotek EPOCH plate reader, Winooski, VT, USA). Intra-assay CV was 4.44% and inter-plate CV was 5.94%.

### 2.5. Statistical analysis

We conducted all statistics in R version 3.6.1 (R-Core-Team, 2019). We examined normality using a Shapiro-Wilk normality test and examined outliers using a Grubbs test in the R package ‘outliers’ (Komsta, 2006). We normalized T using a log scale transformation for all statistical comparisons. There were no differences in testosterone levels between individuals that were collected immediately versus delayed for either sex (t = −0.0081, df = 19.87, p-value = 0.99), so we combined these groups for analysis. To compare T, morphology, and aggression between the sexes at different male breeding stages, we used a one-way ANOVA, followed by a Tukey post hoc test. Comparisons between sexes or male breeding stages were made using Student’s t tests or Wilcoxon tests, depending on normality. To assess the degree of co-variation among T and competitive traits, we used Spearman’s correlations. To control for multiple testing, we used the Benjamini-Hochberg method with the p.adjust function in the R package ‘stats’ (R-Core-Team, 2019).

We also summarized the 5 aggressive behaviors with a principal component analysis (PCA) using the prcomp function. We retained 1 PC with an eigenvalue > 1 (hereafter ‘Aggression PC1’), which explained 64.1% of the variation in the aggressive behaviors (Table 1). Distance loaded negatively onto Aggression PC1, and hover flights, wing spreads, flyovers, and vocalizations loaded positively, such that a more positive PC1 reflects a more aggressive response.

**Table 1.** Loadings for Principal Component Analysis of Aggressive Behavior.

| Vocal Parameter | PC1 |
| --- | --- |
| Eigenvalue | 1.79 |
| Proportion of variance | 64.14% |
| Distance to mount | -0.41 |
| Hover Flights | 0.43 |
| Wing Spreads | 0.46 |
| Flyovers | 0.46 |
| Vocalizations | 0.47 |

## 3. Results

### 3.1. Circulating testosterone varies by sex and male breeding stage

When all males were combined, male and female T were not significantly different (t = −1.58, df = 17.47, p = 0.13). When considering male breeding stage however, the groups were significantly different (F_2,19_ =8.2, p = 0.0027; Fig. 1). Copulating males had significantly higher T levels than incubating males (5.26 ± 2.2 vs 0.54 ± 0.18 ng/mL, Tukey: p = 0.005). Females had T levels (0.51 ± 0.13 ng/mL) similar to incubating males and significantly lower than copulating males (Tukey: p = 0.004). One copulating male had T similar to the average T levels of incubating males, but was not an outlier among other copulating males (p = 0.099); upon collection he was observed to have new feathers growing in over his brood patch.

**Figure 1.**
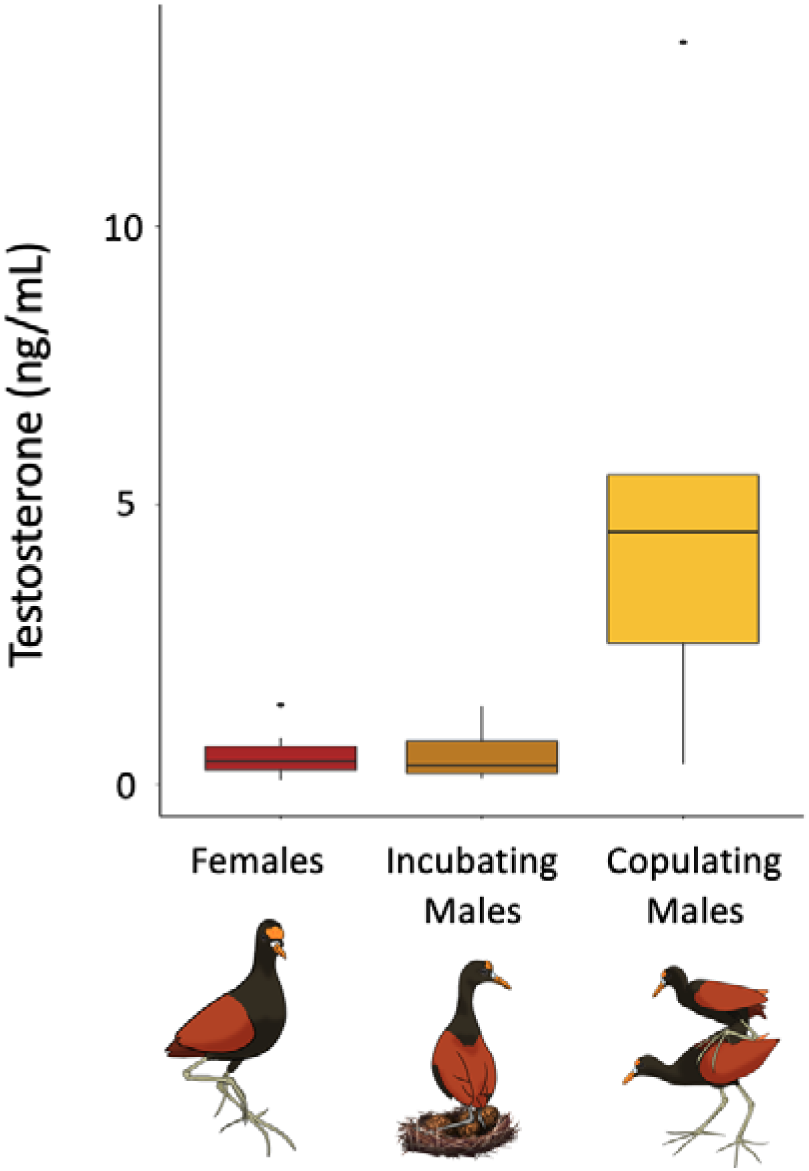
Circulating testosterone levels in female (n = 10) and male jacanas at different breeding stages, incubating (n = 7) and copulating (n = 5). Boxplot horizontal line represents median values outlined by first and third quartiles, with vertical lines representing minimum and maximum values.

### 3.2. Competitive traits vary by sex and male breeding stage

We confirmed female-biased dimorphism in traits putatively involved in competition for mates, as has been previously demonstrated in jacanas (Emlen and Wrege, 2004a; Lipshutz, 2017). Females had significantly longer wing spurs (t = 10.23, df = 20.49, p < 0.0001), larger body mass (t = 21.89, df = 18.54, p < 0.0001), and longer facial shields (t = 9.69, df = 19.0, p < 0.0001) than males in both breeding stages. Between males in different breeding stages, incubating males had significantly lower testis mass than copulating males (t = −2.57, df = 8, p = 0.033), and a trend towards longer wing spurs (t = 2.08, df = 5.15, p = 0.091).

Groups differed significantly in Aggression PC1 (F_2,19_ =3.89, p = 0.038). Males defending their eggs (i.e. incubating males) were marginally more aggressive than copulating males (Tukey: p = 0.089) and significantly more aggressive than females (Tukey: p = 0.049). Aggression did not differ between females and copulating males (Tukey: p = 0.99).

### 3.3. Co-variation between testosterone and competitive traits

In females, T correlated positively with wing spur length, but not with other putatively competitive traits (Table 2). T was positively correlated with testes mass among copulating males, but uncorrelated with any trait among incubating males. T was not correlated with Aggression PC1, regardless of sex or male breeding stage. Individual correlations for each group are plotted in a supplemental figure (Fig. S1).

**Table 2.** Spearman correlations (r_s_) of circulating testosterone with competitive traits for females and males that were incubating or copulating. Significant relationships after Benjamini-Hochberg correction (p ≤ 0.05) are bold.

|  | **Females** | **Incubating Males** | **Copulating Males** |
| --- | --- | --- | --- |
| Wing Spur Length | **$r_s = 0.83$**<br>**$p = 0.036$** | $r_s = -0.07$<br>$p = 0.91$ | $r_s = -0.70$<br>$p = 0.58$ |
| Body Mass | $r_s = 0.09$<br>$p = 0.91$ | $r_s = 0.29$<br>$p = 0.71$ | $r_s = 0.70$<br>$p = 0.58$ |
| Facial Shield Length | $r_s = -0.21$<br>$p = 0.71$ | $r_s = 0.57$<br>$p = 0.58$ | $r_s = 0.4$<br>$p = 0.71$ |
| Gonad Mass | $r_s = 0.25$<br>$p = 0.71$ | $r_s = -0.64$<br>$p = 0.58$ | **$r_s = 0.97$**<br>**$p = 0.036$** |
| Aggression PC1 | $r_s = -0.061$<br>$p = 0.91$ | $r_s = -0.29$<br>$p = 0.71$ | $r_s = 0.45$<br>$p = 0.71$ |

## 4. Discussion

In wild female and male jacanas, we found that differences in T secretion were contingent on male breeding stage, rather than sex alone. Circulating T levels in females were similar to levels observed in males conducting parental care. This is reflective of patterns found in other sex-role reversed species, for which females have similar levels of T in circulation to incubating males, despite female jacanas facing strong mating competition (Emlen and Wrege 2004a). Copulating males had higher T than incubating males, suggesting that T and parental care may constrain one another. Although T varied by sex and male breeding stage in ways that suggest T is shaped by trade-offs between competition and care, we did not find widespread or consistent co-variation between individual differences in T and competitive traits.

In male jacanas, T differs by breeding stage, suggesting that levels of T are shaped by shifting selection pressures from mating effort to parental effort. This pattern is similar to studies of other sex-role reversed species, in which T was similarly low for both females and nesting males, and higher in courting males (Table 3). Stage-related variation in T has been found in non-sex-role reversed species with male parental care (Wingfield et al., 1990). Experimental evidence that exogenous T reduces male parental care in socially monogamous songbirds (Goymann and Davila, 2017; Ketterson et al., 1992; Van Roo, 2004; but see Lynn, 2008) as well as role reversed spotted sandpipers (*Actitis macularia*) (Oring et al., 1989), indicates that high T can inhibit male parental behavior regardless of whether males are the more competitive sex. This idea was supported by a meta-analysis across vertebrates, finding that male parental care predicted T but mating system did not (Hirschenhauser and Oliveira, 2006), and another meta-analysis in humans, finding that fathers have lower T than men without children (Grebe et al., 2019).

**Table 3.** Levels of testosterone in circulation among sex-role reversed females and males in parenting and courting breeding stages.

| Species | Females | Parenting Males | Courting Males | Citation |
| --- | --- | --- | --- | --- |
| African Black Coucal<br>*Centropus grillii* | 0.57 ± 0.06 | 0.59 ± 0.14 | 2.16 ± 0.51 | (Goymann et al., 2004) |
| Wilson's Phalarope<br>*Phalaropus tricolor* | 0.51 ± 0.1 | 0.63 ± 0.15 | 3.61 ± 1.12 | (Fivizzani et al., 1986) |
| Red-necked Phalarope<br>*Phalaropus lobatus* | 0.30 ± 0.19 | 0.35 ± 0.23 | 3.73 ± 2.13 | (Gratto-Trevor et al., 1990) |
| Spotted Sandpiper<br>*Actitis macularia* | 0.18 ± 0.03 | 0.17 ± 0.05 | 0.95 ± 0.41 | (Rissman and Wingfield, 1984) |
| Barred Buttonquail<br>*Turnix suscitator* | 0.2 ± 0.1 | NA | 0.5 ± 0.1 | (Voigt, 2016) |
| Northern Jacana<br>*Jacana spinosa* | 0.51 ± 0.13 | 0.54 ± 0.18 | 5.26 ± 2.2 | this study |

Given the constraints of egg production and parental care typically imposed on females, T levels ought to be less correlated with suites of integrated traits in females than in males (Ketterson et al., 2009). For sex-role reversed species like jacanas, in which males conduct all parental care, we expected this relationship to be reversed between the sexes. However, we did not find strong phenotypic co-variation with T in either sex, except for wing spur length in females and testes mass in copulating males. Considering limited sample sizes and the potential for type II error, we view these analyses as a preliminary but important step towards understanding hormonal regulation of competitive traits and how it may vary based on gonadal sex and parental role. Females with higher T have longer wing spurs, a weapon used to fight over territories and mates. A similar finding in sex-role reversed barred buttonquail females (*Turnix suscitator*) indicated that female but not male levels of T positively correlated with body condition and the size and blackness of the melanin throat patch (Muck and Goymann, 2011). In moorhens, (*Gallinula chloropus)*, the heaviest females, which tend to win most of the competitive interactions, also had higher T levels than lighter females (Eens and Pinxten, 2000). Indeed, females generally have longer wing spurs than males, but do not have higher T than males. In copulating males, only testes mass was correlated with T, a pattern found repeatedly in males of many avian species (Garamszegi et al., 2005). T did not correlate with aggression for either sex, regardless of male breeding stage. This is not unexpected, given that aggression and baseline levels of T do not correlate for many species (Kempenaers et al., 2008; Williams, 2008), although it is possible that such correlations would emerge when T secretion is at its physiological maximum (e.g. endogenous or exogenous activation of the HPG axis). Our observation that incubating males had the lowest T and highest aggression suggests that mechanisms beyond T should be explored in the future (see below). Related to this point, higher aggression in incubating males may serve a parental purpose (i.e. defense of developing young, rather than territorial defense), for which we might expect non-T mechanisms to play a more critical role (Duque-Wilckens and Trainor 2017). Despite these complexities, our analyses nevertheless suggest that there may be sex- and stage-specific variation between T and various competitive traits.

Other endocrine mechanisms are likely to influence the expression of competitive phenotypes; after all, female jacanas have T levels that are on par with incubating males, but they express an exaggerated suite of competitive traits. For example, aggression in role-reversed black coucals is related to progesterone (Goymann et al. 2008). Furthermore, circulating T is only one component of the androgen signaling system, and tissue-specific variation in T production, metabolism, or sensitivity(Ball and Balthazart, 2019; Fuxjager and Schuppe, 2018; Schmidt et al., 2008; Soma, 2006; Staub and De Beer, 1997). Such tissue-specific variability may regulate traits independently of circulating sex steroids (Bentz et al., 2019; Horton et al., 2014; Lipshutz et al., 2019a; Rosvall et al., 2012), and there is some evidence that tissue-specific regulation may be more prevalent for groups in which T levels are depressed (Demas et al., 2007; Rosvall, 2013b). Sex differences in androgenic signaling in the brain have been found in several sex-role reversed species; female black coucals and barred button quails had higher mRNA expression of androgen receptors (AR) in neural regions implicated in the control of aggressive and sexual behavior (Voigt, 2016; Voigt and Goymann, 2007). An exciting next step is to use the unique jacana system to explore how gonadal sex and parental role influence tissue-level regulation of competitive phenotypes.

As tropical birds that are also sex role-reversed, the jacana system sets up a unique situation in which females experience nearly year-round selection in relation to mating competition. Territorial female jacanas breed simultaneously with multiple males in their harems, and males copulate and incubate asynchronously (Emlen and Wrege, 2004a). All females in this study displayed clear signs of fertility (i.e. hierarchical follicles), whereas males had large testes only when copulating, but small testes when incubating. In this sense, the continuously fertile gonadal state of female jacanas is similar to other species for which males maintain large, fertile testes throughout female incubation, demonstrating a direct reversal of sex differences in fertility seen in non role-reversed systems. Likewise, the cycling gonadal size of male jacanas from copulation to incubation is analogous to other species for which female ovaries return to a non-fertile state during incubation (Williams, 2012). The essentially continual state of female jacana fertility, paired with sex-role reversal, should generate stronger or more uniform selection on competitive traits, which are advantageous during most of the year and do not experience counter-selection in relation to wintering or migratory states. Our examination of T and competitive traits in this classic system of sex role reversal brings us one step closer to understanding how gonadal sex and parental roles influence mechanisms of behavior.

## Acknowledgments

This material is supported by IU Biology, a National Science Foundation Graduate Research Fellowship Grant No. 1154145, Doctoral Dissertation Improvement Grant IOS-1818235, and a Smithsonian Tropical Research Institute short-term fellowship to SEL. Any opinion, findings, and conclusions or recommendations expressed in this material are those of the authors(s) and do not necessarily reflect the views of the National Science Foundation. Scientific collection in Panama is done with permission from landowners and prior approval of MiAmbiente, Panama’s environmental authority (permit number: SE/A-17-18), and the Institutional Animal Care and Use Committee of the Smithsonian Tropical Research Institute (IACUC permit: 2018-0116-2021) and the University of Tennessee (IACUC permit: 2573). We thank the Rosvall lab for feedback, Evan Buck for assistance in the field, Matt Fuxjager and Meredith Miles for assistance importing and storing samples, the STRI Bird Collection for preparing taxidermic mounts, Mae Berlow for the jacana illustrations, and Elizabeth Derryberry for conceptual design of the study.

## Literature Cited

1. Ball, G.F., Balthazart, J., 2019. Hormones and Behavior The neuroendocrine integration of environmental information, the regulation and action of testosterone and the challenge hypothesis. Horm. Behav. 104574. https://doi.org/10.1016/j.yhbeh.2019.104574

2. Bentz, A.B., Dossey, E.K., Rosvall, K.A., 2019. Tissue-specific gene regulation corresponds with seasonal plasticity in female testosterone. Gen. Comp. Endocrinol. 270, 26–34. https://doi.org/10.1016/j.ygcen.2018.10.001

3. Cain, K.E., Ketterson, E.D., 2012. Competitive females are successful females; phenotype, mechanism, and selection in a common songbird. Behav. Ecol. Sociobiol. 66, 241–252. https://doi.org/10.1007/s00265-011-1272-5

4. Clutton-Brock, T., 2009. Sexual selection in females. Anim. Behav. 77, 3–11. https://doi.org/10.1016/j.anbehav.2008.08.026

5. Cox, R.M., Stenquist, D.S., Calsbeek, R., 2009. Testosterone, growth and the evolution of sexual size dimorphism. J. Evol. Biol. 22, 1586–1598. https://doi.org/10.1111/j.1420-9101.2009.01772.x

6. Demas, G.E., Cooper, M.A., Albers, H.E., Soma, K.K., 2007. Novel Mechanisms Underlying Neuroendocrine Regulation of Aggression: A Synthesis of Rodent, Avian, and Primate Studies, in: Lajtha, A., Blaustein, J.D. (Eds.),. Springer US, Boston, MA, pp. 337–372. https://doi.org/10.1007/978-0-387-30405-2_8

7. DeVries, M.S., Winters, C.P., Jawor, J.M., 2012. Testosterone elevation and response to gonadotropin-releasing hormone challenge by male northern cardinals (Cardinalis cardinalis) following aggressive behavior. Horm. Behav. 62, 99–105. https://doi.org/10.1016/j.yhbeh.2012.05.00

8. Duque-Wilckens, N., Trainor, B.C., 2017. Behavioral neuroendorinology of female aggression. Oxford Research Encyclopedia of Neuroscience. 10.1093/acrefore/9780190264086.013.11

9. Eens, M., Pinxten, R., 2000. Sex-role reversal in vertebrates: Behavioural and endocrinological accounts. Behav. Processes 51, 135–147. https://doi.org/10.1016/S0376-6357(00)00124-8

10. Eens, M., Van Duyse, E., Berghman, L., Pinxten, R., 2000. Shield characteristics are testosterone-dependent in both male and female moorhens. Horm. Behav. 37, 126–134. https://doi.org/10.1006/hbeh.1999.1569

11. Emlen, S.T., Demong, N.J., Emlen, D.J., 1989. Experimental induction of infanticide in female Wattled Jacanas. Auk 41, 225–230.

12. Emlen, S.T., Oring, L.W., 1977. Ecology, sexual selection, and evolution of mating systems. Science (80-.). 197, 215–223.

13. Emlen, S.T., Wrege, P.H., 2004a. Size dimorphism, intrasexual competition, and sexual selection in Wattled Jacana (*Jacana Jacana*), a sex-role-reversed shorebird in Panama. Auk 121, 391–403. https://doi.org/10.1642/0004-8038(2004)121[0391:SDICAS]2.0.CO;2

14. Emlen, S.T., Wrege, P.H., 2004b. Division of labour in parental care behaviour of a sex-role-reversed shorebird, the Wattled Jacana. Anim. Behav. 68, 847–855. https://doi.org/10.1016/j.anbehav.2003.08.034

15. Fivizzani, A.J., Colwell, M.A., Oring, L.W., 1986. Plasma steroid hormone levels in free-living Wilson’s phalaropes, Phalaropus tricolor. Gen. Comp. Endocrinol. 62, 137–144. https://doi.org/https://doi.org/10.1016/0016-6480(86)90102-4

16. Fuxjager, M.J., Schuppe, E.R., 2018. Androgenic signaling systems and their role in behavioral evolution. J. Steroid Biochem. Mol. Biol. 184, 47–56. https://doi.org/10.1016/j.jsbmb.2018.06.004

17. Garamszegi, L., Hirschenhauser, K., Bókony, V., Eens, M., Hurtrez-Boussès, S., Møller, A., Oliveira, R., Wingfield, J., 2008. Latitudinal Distribution, Migration, and Testosterone Levels in Birds. Am. Nat. 172, 533–546. https://doi.org/10.1086/590955

18. Garamszegi, L.Z., Eens, M., Hurtrez-Boussès, S., Møller, A.P., 2005. Testosterone, testes size, and mating success in birds: a comparative study. Horm. Behav. 47, 389–409. https://doi.org/https://doi.org/10.1016/j.yhbeh.2004.11.008

19. George, E.M., Rosvall, K.A., 2018. Testosterone production and social environment vary with breeding stage in a competitive female songbird. Horm. Behav. 103, 28–35. https://doi.org/10.1016/j.yhbeh.2018.05.015

20. Goymann, W., Davila, P.F., 2017. Acute peaks of testosterone suppress paternal care: evidence from individual hormonal reaction norms. Proc. R. Soc. B 284, 20170632.

21. Goymann, W., Wingfield, J.C., 2014. Male-to-female testosterone ratios, dimorphism, and life history - What does it really tell us? Behav. Ecol. 25, 685–699. https://doi.org/10.1093/beheco/aru019

22. Goymann, W., Wittenzellner, A., Schwabl, I., Makomba, M., 2008. Progesterone modulates aggression in sex-role reversed female African black coucals. Proc. R. Soc. B Biol. Sci. 275, 1053–1060. https://doi.org/10.1098/rspb.2007.1707

23. Goymann, W., Wittenzellner, A., Wingfield, J.C., 2004. Competing females and caring males. Polyandry and sex-role reversal in African black coucals, Centropus grillii. Ethology 110, 807–823. https://doi.org/10.1111/j.1439-0310.2004.01015.x

24. Gratto-Trevor, C.L., Fivizzani, A.J., Oring, L.W., Cooke, F., 1990. Seasonal changes in gonadal steroids of a monogamous versus a polyandrous shorebird. Gen. Comp. Endocrinol. 80, 407–418. https://doi.org/10.1016/0016-6480(90)90190-W

25. Grebe, N.M., Sarafin, R.E., Strenth, C.R., Zilioli, S., 2019. Pair-bonding, fatherhood, and the role of testosterone: A meta-analytic review. Neurosci. Biobehav. Rev. 98, 221–233. https://doi.org/https://doi.org/10.1016/j.neubiorev.2019.01.010

26. Hare, R.M., Simmons, L.W., 2018. Sexual selection and its evolutionary consequences in female animals. Biol. Rev. https://doi.org/10.1111/brv.12484

27. Hau, M., Ricklefs, R.E., Wikelski, M., Lee, K.A., Brawn, J.D., 2010. Corticosterone, testosterone and life-history strategies of birds. Proc. R. Soc. B 3203–3212. https://doi.org/10.1098/rspb.2010.0673

28. Hirschenhauser, K., Oliveira, R.F., 2006. Social modulation of androgens in male vertebrates: Meta-analyses of the challenge hypothesis. Anim. Behav. 71, 265–277. https://doi.org/10.1016/j.anbehav.2005.04.014

29. Höhn, E.O., 1970. Gonadal hormone concentrations in northern phalaropes in relation to nuptial plumage. Can. J. Zool. 48, 400–401. https://doi.org/10.1139/z70-067

30. Höhn, E.O., Cheng, S.C., 1967. Gonadal hormones in Wilson’s phalarope (Steganopus tricolor) and other birds in relation to plumage and sex behavior. Gen. Comp. Endocrinol. 8, 1–11. https://doi.org/https://doi.org/10.1016/0016-6480(67)90105-0

31. Horton, B.M., Hudson, W.H., Ortlund, E.A., Shirk, S., Thomas, J.W., Young, E.R., Zinzow-Kramer, W.M., Maney, D.L., 2014. Estrogen receptor α polymorphism in a species with alternative behavioral phenotypes. Proc. Natl. Acad. Sci. 111, 1443–1448. https://doi.org/10.1073/pnas.1317165111

32. Jawor, J.M., Mcglothlin, J.W., Casto, J.M., Greives, T.J., Snajdr, E.A., Bentley, G.E., Ketterson, E.D., 2007. Testosterone response to GnRH in a female songbird varies with stage of reproduction: implications for adult behavior and maternal effects. Funct. Ecol. 21, 767–775. https://doi.org/10.1111/j.1365-2435.2007.01280.x

33. Jenni, D.A., Collier, G., 1972. Polyandry in the American Jaçana. Auk 89(4), 743–765.

34. Johnson, D.D.P., Briskie, J. V., 1999. Sperm Competition and Sperm Length in Shorebirds. Condor 101, 848–854.

35. Kempenaers, B., Peters, A., Foerster, K., 2008. Sources of individual variation in plasma testosterone levels. Philos. Trans. R. Soc. B Biol. Sci. 363, 1711–1723. https://doi.org/10.1098/rstb.2007.0001

36. Ketterson, E.D., Atwell, J.W., McGlothlin, J.W., 2009. Phenotypic integration and independence: Hormones, performance, and response to environmental change. Integr. Comp. Biol. 49, 365–379. https://doi.org/10.1093/icb/icp057

37. Ketterson, E.D., Nolan, V., Wolf, L., Ziegenfus, C., 1992. Testosterone and Avian Life Histories: Effects of Experimentally Elevated Testosterone on Behavior and Correlates of Fitness in the Dark-Eyed Junco (Junco hyemalis). Am. Nat. 140, 980–999. https://doi.org/10.1086/285451

38. Komsta, L., 2006. Processing data for outliers. R News 6, 21–26.

39. Lipshutz, S.E., 2017. Divergent competitive phenotypes between females of two sex-role-reversed species. Behav. Ecol. Sociobiol. 71. https://doi.org/10.1007/s00265-017-2334-0

40. Lipshutz, S.E., George, E.M., Bentz, A.B., Rosvall, K.A., 2019a. Molecular and Cellular Endocrinology Evaluating testosterone as a phenotypic integrator: From tissues to individuals to species. Mol. Cell. Endocrinol. 496, 110531. https://doi.org/10.1016/j.mce.2019.110531

41. Lipshutz, S.E., Seehausen, O., Meier, J.I., Derryberry, G.E., Miller, M.J., Derryberry, E.P., 2019b. Differential introgression of a female competitive trait in a hybrid zone between sex-role reversed species. Evolution (N. Y). 73, 188–201. https://doi.org/10.1111/evo.13675

42. Lynn, S.E., 2008. Behavioral insensitivity to testosterone: why and how does testosterone alter paternal and aggressive behavior in some avian species but not others? Gen. Comp. Endocrinol. 157, 233–240. https://doi.org/10.1016/j.ygcen.2008.05.009

43. Malo, A.F., Roldan, E.R.S., Garde, J.J., Soler, A.J., Vicente, J., Gortazar, C., Gomendio, M., 2009. What does testosterone do for red deer males□? Proc. R. Soc. B Biol. Sci. 1, 971–980. https://doi.org/10.1098/rspb.2008.1367

44. McGlothlin, J.W., Jawor, J.M., Greives, T.J., Casto, J.M., Phillips, J.L., Ketterson, E.D., 2008. Hormones and honest signals: males with larger ornaments elevate testosterone more when challenged. J. Evol. Biol. 21, 39–48. https://doi.org/10.1111/j.1420-9101.2007.01471.x

45. Muck, C., Goymann, W., 2011. Throat patch size and darkness covaries with testosterone in females of a sex-role reversed species. Behav. Ecol. 20. https://doi.org/10.1093/beheco/arr133

46. Oring, L.W., Fivizzani, A.J., El Halawani, M.E., 1989. Testosterone-Induced Inhibition of Incubation in the Spotted Sandpiper (Actitis macularia). Horm. Behav. 23, 412–423.

47. Preston, B.T., Stevenson, I.R., Lincoln, G.A., Monfort, S.L., Pilkington, J.G., Wilson, K., 2012. Testes size, testosterone production and reproductive behaviour in a natural mammalian mating system. J. Anim. Ecol. 81, 296–305. https://doi.org/10.1111/j.1365-2656.2011.01907.x

48. R-Core-Team, 2019. R: A language and environment for statistical computing.

49. Rissman, E.F., Wingfield, J.C., 1984. Hormonal correlates of polyandry in the spotted sandpiper, Actitis macularia. Gen. Comp. Endocrinol. 56, 401–405. https://doi.org/10.1016/0016-6480(84)90082-0

50. Rosvall, K.A., 2013a. Proximate perspectives on the evolution of female aggression: good for the gander, good for the goose? Philos. Trans. R. Soc. B Biol. Sci. 368, 20130083. https://doi.org/10.1098/rstb.2013.0083

51. Rosvall, K.A., 2013b. Life History Trade-Offs and Behavioral Sensitivity to Testosterone: An Experimental Test When Female Aggression and Maternal Care Co-Occur. PLoS One 8. https://doi.org/10.1371/journal.pone.0054120

52. Rosvall, K.A., Bentz, A.B., George, E.M., 2019. How research on female vertebrates contributes to an expanded challenge hypothesis. Horm. Behav. 104565. https://doi.org/https://doi.org/10.1016/j.yhbeh.2019.104565

53. Rosvall, K.A., Bergeon Burns, C.M., Barske, J., Goodson, J.L., Schlinger, B.A., Sengelaub, D.R., Ketterson, E.D., 2012. Neural sensitivity to sex steroids predicts individual differences in aggression: implications for behavioural evolution. Proc. R. Soc. B Biol. Sci. 279, 3547–3555. https://doi.org/10.1098/rspb.2012.0442

54. Schmidt, K.L., Pradhan, D.S., Shah, A.H., Charlier, T.D., Chin, E.H., Soma, K.K., 2008. Neurosteroids, immunosteroids, and the Balkanization of endocrinology. Gen. Comp. Endocrinol. 157, 266–274. https://doi.org/10.1016/j.ygcen.2008.03.025

55. Soma, K.K., 2006. Testosterone and Aggression: Berthold, Birds and Beyond. J. Neuroendocrinol. 18, 543–551. https://doi.org/10.1111/j.1365-2826.2006.01440.x

56. Staub, N.L., De Beer, M., 1997. The role of androgens in female vertebrates. Gen. Comp. Endocrinol. 108, 1–24. https://doi.org/10.1006/gcen.1997.6962

57. Stephens, M.L., 1984. Aggressive behavior of the polyandrous Northern Jacana (*Jacana spinosa*). Auk 101, 508–518.

58. Tobias, J.A., Montgomerie, R., Lyon, B.E., 2012. The evolution of female ornaments and weaponry: social selection, sexual selection and ecological competition. Philos. Trans. R. Soc. B Biol. Sci. 367, 2274–2293. https://doi.org/10.1098/rstb.2011.0280

59. Van Roo, B.L., 2004. Exogenous testosterone inhibits several forms of male parental behavior and stimulates song in a monogamous songbird: The blue-headed vireo (Vireo solitarius). Horm. Behav. 46, 678–683. https://doi.org/10.1016/j.yhbeh.2004.06.011

60. Voigt, C., 2016. Neuroendocrine correlates of sex-role reversal in barred buttonquails. Proc. R. Soc. B Biol. Sci. 283. https://doi.org/10.1098/rspb.2016.1969

61. Voigt, C., Goymann, W., 2007. Sex-role reversal is reflected in the brain of African black coucals (*Centropus grillii*). Dev. Neurobiol. 67, 1560–1573. https://doi.org/10.1002/dneu.20528

62. Williams, T.D., 2012. Physiological adaptations for breeding in birds. Princeton University Press.

63. Williams, T.D., 2008. Individual variation in endocrine systems: moving beyond the ‘tyranny of the Golden Mean.’ Philos. Trans. R. Soc. B Biol. Sci. 363, 1687–1698. https://doi.org/10.1098/rstb.2007.000

64. Wingfield, J.C., Ramenofsky, M., Hegner, R.E., Ball, B.F. in press. Wither the challenge hypothesis? Horm. Behav. https://doi.org/10.1016/j.yhbeh.2019.104588

65. Wingfield, J.C., Hegner, R.E., Dufty, A.M., Ball, G.F., 1990. The “Challenge Hypothesis”: Theoretical Implications for Patterns of Testosterone Secretion, Mating Systems, and Breeding Strategies. Am. Nat. 136, 829–846.

66. Wingfield, J.C., Wada, M., 1989. Changes in plasma levels of testosterone during male-male interactions in the song sparrow, Melospiza melodia: time course and specificity of response. J. Comp. Physiol. A 166, 189–194. https://doi.org/10.1007/BF00193463

